# Glycomics Profiling Reveals a Stromal Chondroitin-4-sulfate State that Shapes Platinum Response in Ovarian Cancer

**DOI:** 10.1101/2025.10.06.675352

**Authors:** Erica J. Peterson, James D. Hampton, Thomas M. Clausen, Ava R.S. Beaudin, Sharanya P. Deshmukh, Mikhail G. Dozmorov, Joseph B. McGee Turner, Amrita Basu, Elena Ethel Vidal-Calvo, Corinne E. Leonard, Priscilla Y. Hwang, Mads Daugaard, Ali Salanti, Ryan J. Weiss, Jennifer E. Koblinski, Larisa Litovchick, Nicholas P. Farrell

**Affiliations:** Division of Hematology, Oncology, and Palliative Care, Department of Internal Medicine, Virginia Commonwealth University; Richmond, VA, USA; Massey Comprehensive Cancer Center, Virginia Commonwealth University; Richmond, VA, USA; Bioplatinum Technologies LLC; Richmond, VA, USA; Department of Chemistry, Virginia Commonwealth University; Richmond, VA, USA; Center for translational Medicine, Department of Immunology and Microbiology, University of Copenhagen, Copenhagen, Denmark; VAR2 Pharmaceuticals ApS, Copenhagen, Denmark; Department of Biochemistry and Molecular Biology, University of Georgia; Athens, GA, USA; Complex Carbohydrate Research Center, University of Georgia; Athens, GA, USA; Department of Biomedical Engineering, Virginia Commonwealth University; Richmond, VA, USA; Department of Urologic Sciences, Faculty of Medicine, University of British Columbia, Vancouver, BC, Canada V5Z 1M9; Vancouver Prostate Centre, Vancouver, BC, Canada V6H 3Z6; Department of Pathology, Virginia Commonwealth University; Richmond, VA, USA

## Abstract

For women with advanced ovarian cancer (OC), remission is typically achieved through surgery and platinum-based combination chemotherapy, but the duration of response varies widely, and reliable biomarkers of platinum sensitivity remain limited. Because tumor-associated glycosaminoglycans (GAGs) influence extracellular signaling and drug interactions, but are not directly resolved by conventional transcriptomic approaches, we investigated whether direct glycomics profiling could identify a functionally relevant determinant of platinum response. Glycan reductive isotope labeling-LC/MS of ovarian cancer patient-derived xenografts (PDXs) showed that carboplatin-resistant tumors were enriched for chondroitin-4-sulfate (C4S). Single-cell transcriptomic analyses localized candidate proteoglycan scaffolds, including VCAN and BGN, primarily to stromal fibroblast compartments, supporting a stromal matrix source while underscoring that RNA measurements cannot define mature GAG abundance or fine structure. Analysis of tissue microarrays representing multiple ovarian cancer histotypes revealed high C4S expression, including in subtypes with intrinsic platinum resistance. CRISPR-mediated double *XYLT1* and *XYLT2* knockout reduced C4S levels, slowed ES2 tumor growth, reduced spheroid invasion, altered intraperitoneal dissemination, and eliminated ascites in the ES2 model. *XYLT1*/*2* knockout also shifted platinum pharmacology: carboplatin uptake and DNA platination increased, whereas Triplatin accumulation and activity decreased. In PDX models, tissue rVAR2-V5 staining correlated with direct glycomics measurements and identified carboplatin-resistant tumors selectively sensitive to the GAG-targeting platinum agent Triplatin. These findings nominate a stromal-associated C4S-rich extracellular state as a non-genetic determinant of platinum response and tumor behavior, and support tissue-level C4S assessment for biomarker-guided evaluation of Triplatin.

**One Sentence Summary:** A stromal C4S-rich glycan state is associated with carboplatin resistance and Triplatin sensitivity in ovarian cancer.

## INTRODUCTION

Ovarian cancer remains one of the most lethal gynecologic malignancies and a leading cause of cancer-related death in women *(1)*. Standard treatment for advanced disease relies on cytoreductive surgery followed by combination chemotherapy with paclitaxel and a platinum-based agent, most commonly carboplatin. Although most patients initially respond, relapse is common and recurrent tumors frequently become refractory to subsequent platinum treatment*(2)*. Platinum-resistant ovarian cancer, defined clinically by recurrence within six months of platinum therapy, is associated with poor outcomes, and some histologic subtypes, including clear cell and low-grade serous carcinomas, show intrinsic resistance to standard platinum regimens *(3–5)*. These clinical challenges require biomarkers that identify tumors in which carboplatin benefit may be limited and that nominate rational therapeutic alternatives.

Most biomarker efforts for platinum response have focused on genomic, transcriptomic, or protein-based determinants such as altered transporter expression, DNA repair capacity, and apoptotic signaling *(6, 7)*. These approaches do not capture the extracellular glycan environment that can influence drug distribution at the tumor-stroma interface.

Glycosaminoglycans (GAGs) are highly sulfated, linear polysaccharides displayed on proteoglycans at the cell surface and in the extracellular matrix, where they shape growth factor signaling, cell adhesion, angiogenesis, and immune interactions *(8–18)*. GAG biosynthesis is non-template-driven and depends on coordinated chain initiation, elongation, and sulfation by multiple enzymes. As a result, transcript abundance of individual proteoglycans or biosynthetic enzymes can only partially infer the abundance, chain length, or fine structure of the mature GAGs that mediate biological activity (Fig. 1).

**Fig. 1.**
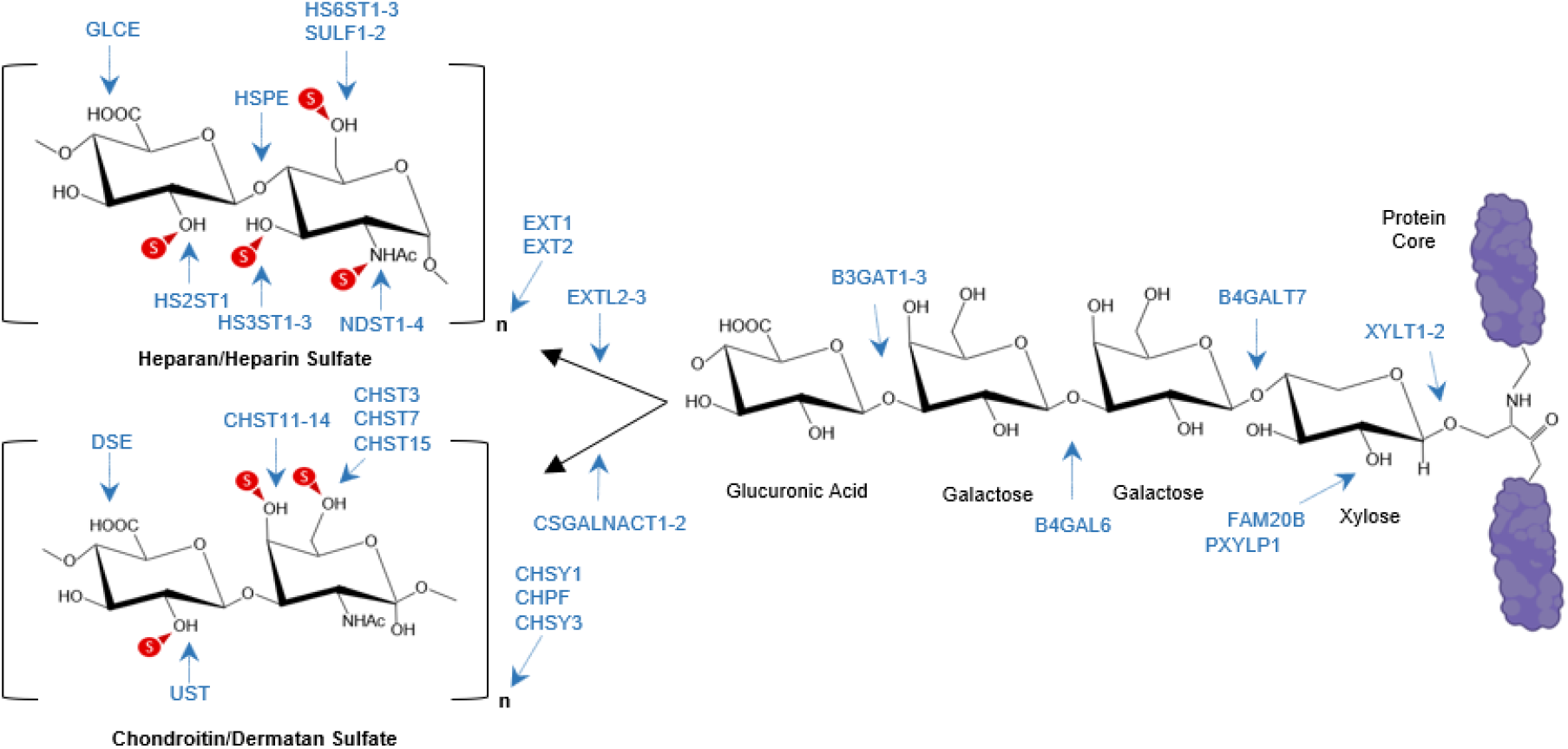
GAG levels are determined by expression of many enzymes. Heparan sulfate or chondroitin /dermatan sulfate attachment sites in a proteoglycan protein core with a linker tetrasaccharide. Biosynthetic steps associated with the listed genes are indicated by arrows.

Among tumor-associated GAGs, chondroitin-4-sulfate and oncofetal chondroitin sulfate structures (C4S/ofCS) are of particular interest because they are enriched across many cancers and show limited expression in normal adult tissues outside the placenta *(15, 19–28)*. Prior work showed that the cationic trinuclear platinum compound, Triplatin, interacts with sulfated GAGs and that depletion of cell-surface GAGs reduces its cellular accumulation and activity *(29–31)*. Whether tumor-associated GAG architecture also modifies carboplatin distribution and response has remained unresolved. Here, direct glycomics profiling, tissue-level staining, and GAG-biosynthesis perturbation identify a stromal-associated C4S/ofCS-rich state linked to carboplatin resistance and Triplatin sensitivity. These results support direct glycomics or tissue-level assays, rather than transcriptomic inference alone, as translational readouts of an extracellular determinant of platinum response in ovarian cancer.

## RESULTS

### Subhead 1: Chondroitin sulfate enrichment is associated with carboplatin resistance

To determine whether tumor-associated GAGs are linked to platinum response, we profiled high-grade serous ovarian cancer PDX tumors using glycan reductive isotope labeling-LC/MS (GRIL-LC/MS)*(32–34)*. Across the initial discovery cohort (Table 1), chondroitin sulfate (CS) was the dominant GAG class, and chondroitin-4-sulfate was the predominant monosulfated CS disaccharide species (Fig. 2A-C and fi g. S1).

**Table 1.**
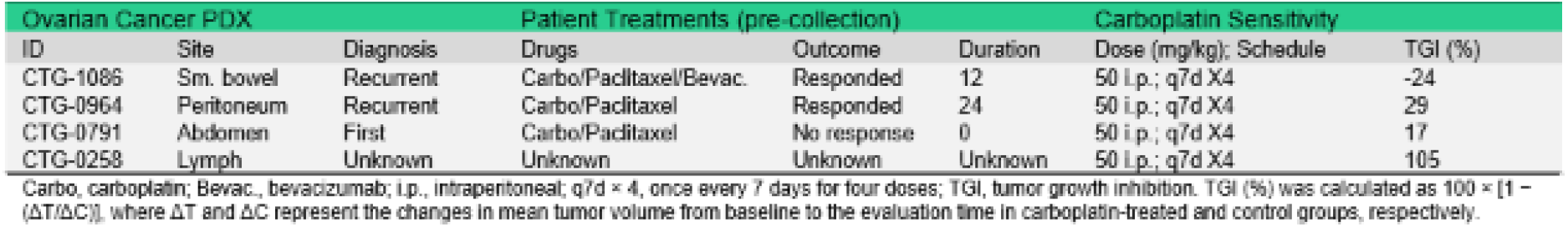
PDX characteristics, patient treatments, and platinum sensitivity. Carbo, carboplatin; Bevac., bevacizumab; i.p., intraperitoneal; q?d × 4, once every 7 days for four doses; TGI, tumor growth inhibition. TGI (%) was calculated as 100 × [1 - (ΔT/ΔC)], where ΔT and ΔC represent the changes in mean tumor volume from baseline to the evaluation time in carboplatin-treated and control groups, respectively.

**Fig. 2.**
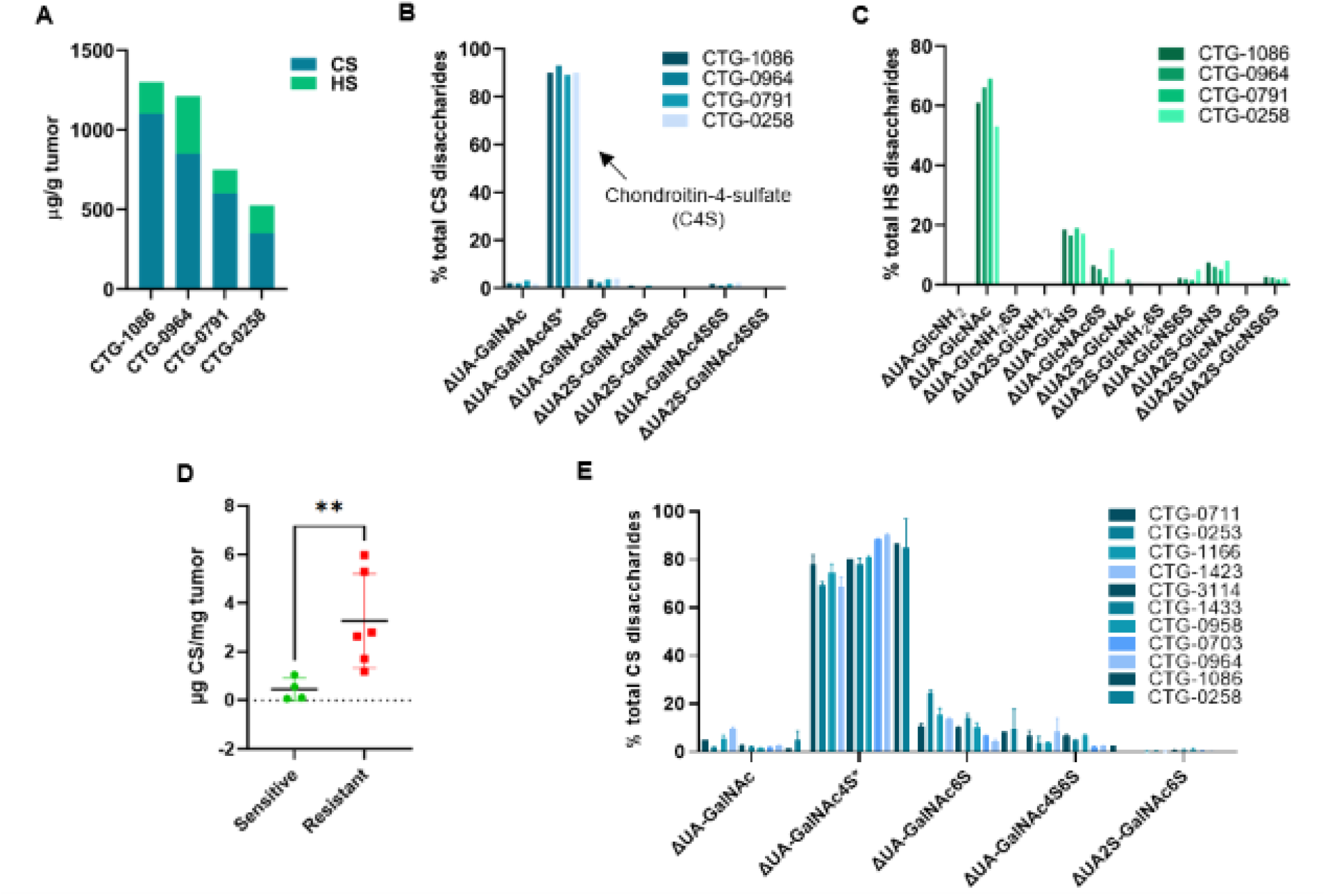
Carboplatin resistance correlates with chondroitin-4-sulfate levels in OC PDX models. **(A)** Quantitative analysis of total GAG levels in OC PDXs using GRIL-MS **(B)** Compositional analysis of chondroitin sulfate in OC PDX using GRIL-MS **(C)** Compositional analysis of heparan sulfate in OC PDX using GRIL-MS **(D)** Quantitative analysis of the total GAG levels measured in 11 PDX models (2 tumors per model). PDX models were determined to be carboplatin-sensitive or resistant if the tumor growth inhibition (TGI) was >50% or <50%, respectively, in mice treated with 50 mg/kg i.p. carboplatin q7dx4. Tumor samples and drug screens provided by Champions Oncology. ^**^p<0.01, Student’s t-test. **(E)** Compositional analysis of chondroitin sulfate in OC PDX using GRIL-MS.

Bulk RNA-seq analysis of genes encoding selected proteoglycan core proteins, chondroitin sulfate initiation and elongation enzymes, and sulfotransferases revealed expression patterns broadly concordant with the glycomics profile; however, transcript abundance could not directly define the abundance or fine structure of mature GAGs (fig. S2). In an expanded panel of ovarian cancer PDX models with known carboplatin response, carboplatin-resistant tumors contained significantly higher C4S than carboplatin-sensitive tumors (Fig. 2D,E). These data identify a reproducible glycomics phenotype linked to platinum response and establish that the relevant variable is not simply proteoglycan transcript abundance, but the abundance and composition of the mature GAG chains themselves.

### Subhead 2: Fibroblasts express candidate proteoglycan scaffolds

To identify cellular compartments that could generate the matrix associated with the C4S-rich state, we analyzed public single-cell RNA-seq datasets spanning normal ovary, primary ovarian or fallopian tube tumors, and metastatic abdominal and colonic sites (Fig. 3A)*(35–37)*.

**Fig. 3.**
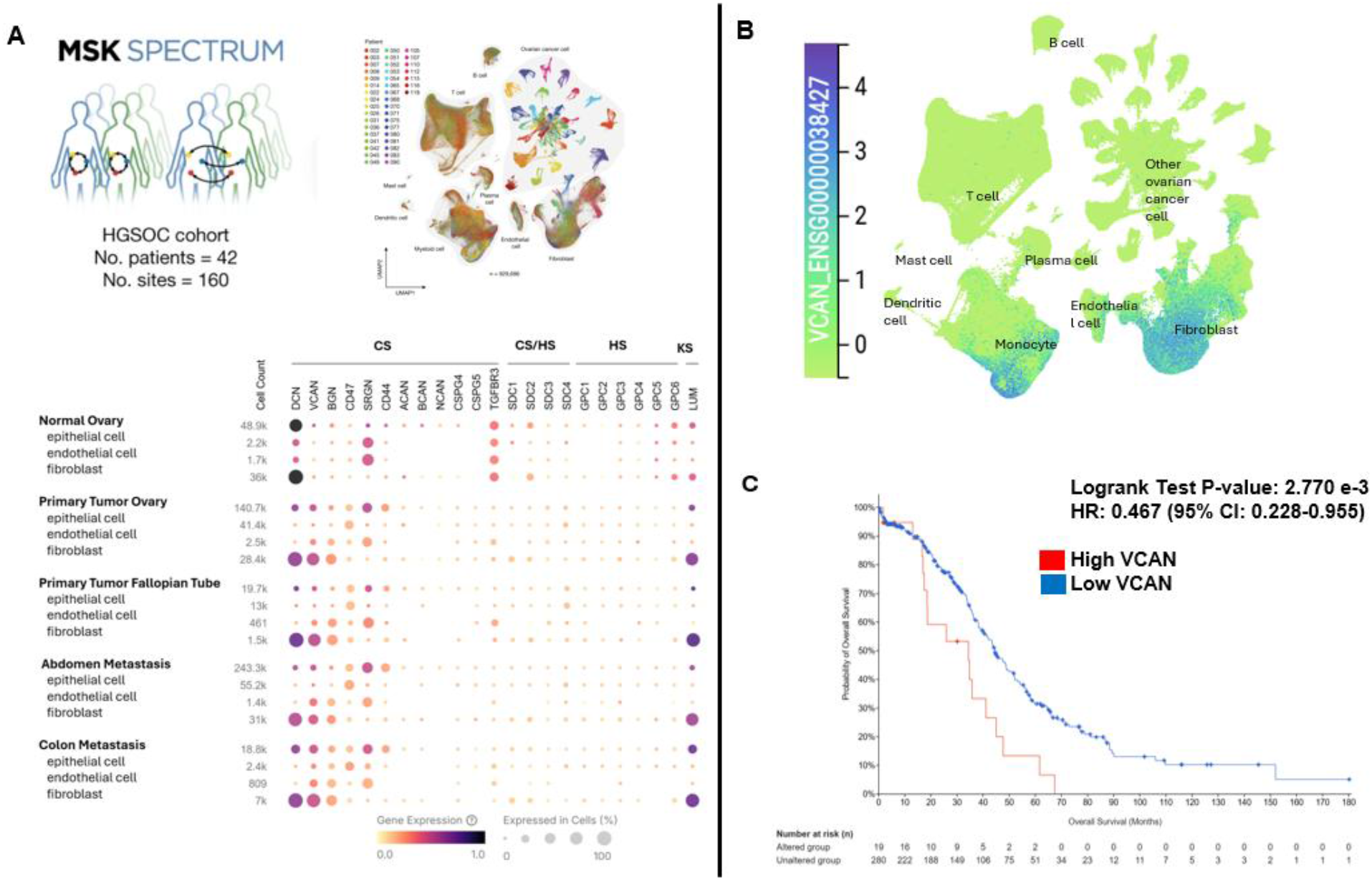
Fibroblast-enriched proteoglycan expression is associated with worse survival in high-grade serous ovarian cancer. **(A)** Single-cell RNAseq analysis of proteoglycan genes in normal ovarian tissue, primary ovarian/fallopian tube tumor and abdominal and colon metastasis. Multi-site tissue samples were collected from patients undergoing pre-treatment laparoscopy or primary debulking surgeries [Dataset (https://cellxgene.cziscience.com/collections/e5f58829-1a66-40b5-a624-9046778e74f5)]. Normal ovary tissue analysis comes from the first-draft human cell atlas of over 1.1M cells from 28 organs of 24 normal human subjects [Dataset (https://cellxgene.cziscience.com/collections/e5f58829-1a66-40b5-a624-9046778e74f5)]. **(B)** VCAN expression across multiple cell types. **(C)** cBioportal analysis showing survival time of patients with high and low levels of VCAN expression. Dataset: Ovarian Serous Cystadenocarcinoma (TCGA PanCancer Atlas); mRNA expression z-scores relative to diploid samples (RNA Seq VS RSEM). z-score threshold +/-1.5

Expression of candidate proteoglycan core proteins, including VCAN and BGN, was enriched predominantly in stromal fibroblast populations, with additional contributions from immune-associated cell types (Fig. 3B). In independent transcriptomic datasets, higher expression of VCAN and CS-associated genes (HSPG2, CSGALNACT2, CHST15, CHSY3 and BCAN) were associated with reduced survival times and shorter progression free intervals after treatment with chemotherapy (Fig. 3C, S3). Together, these data nominate stromal fibroblasts and tumor-associated matrix programs as likely sources of proteoglycan scaffolds associated with the C4S-rich state.

### Subhead 3: C4S/ofCS is prevalent in resistant histotypes

To estimate the clinical prevalence of the C4S-rich state, we stained ovarian cancer tissue microarrays (TMAs) with rVAR2-V5, a recombinant malarial-derived peptide that binds the tumor-associated C4S/ofCS motif and quantified the proportion score (PS) by digital pathology. In the multi-histology OvCa2 cohort, high C4S/ofCS expression was common across epithelial ovarian cancer subtypes and was especially prevalent in low-grade serous carcinoma and clear cell carcinoma, both of which are recognized for poor responses to standard platinum therapy (Fig. 4A-D)(*(4, 5)*. A clear cell carcinoma tissue TMA showed similarly high prevalence of elevated staining (Fig. 4E). Consistent with the single-cell analyses, C4S/ofCS signal frequently colocalized with stromal-rich regions (fig. S4), supporting the concept that stromal fibroblast-associated matrix programs are a major source of the biomarker-positive state. Taken together, these data extend the discovery and mechanistic findings into human tissue and support direct tissue-level measurement of C4S/ofCS as a translational strat egy for identifying ovarian tumors in which carboplatin efficacy may be limited and Triplatin may be favored.

**Fig. 4.**
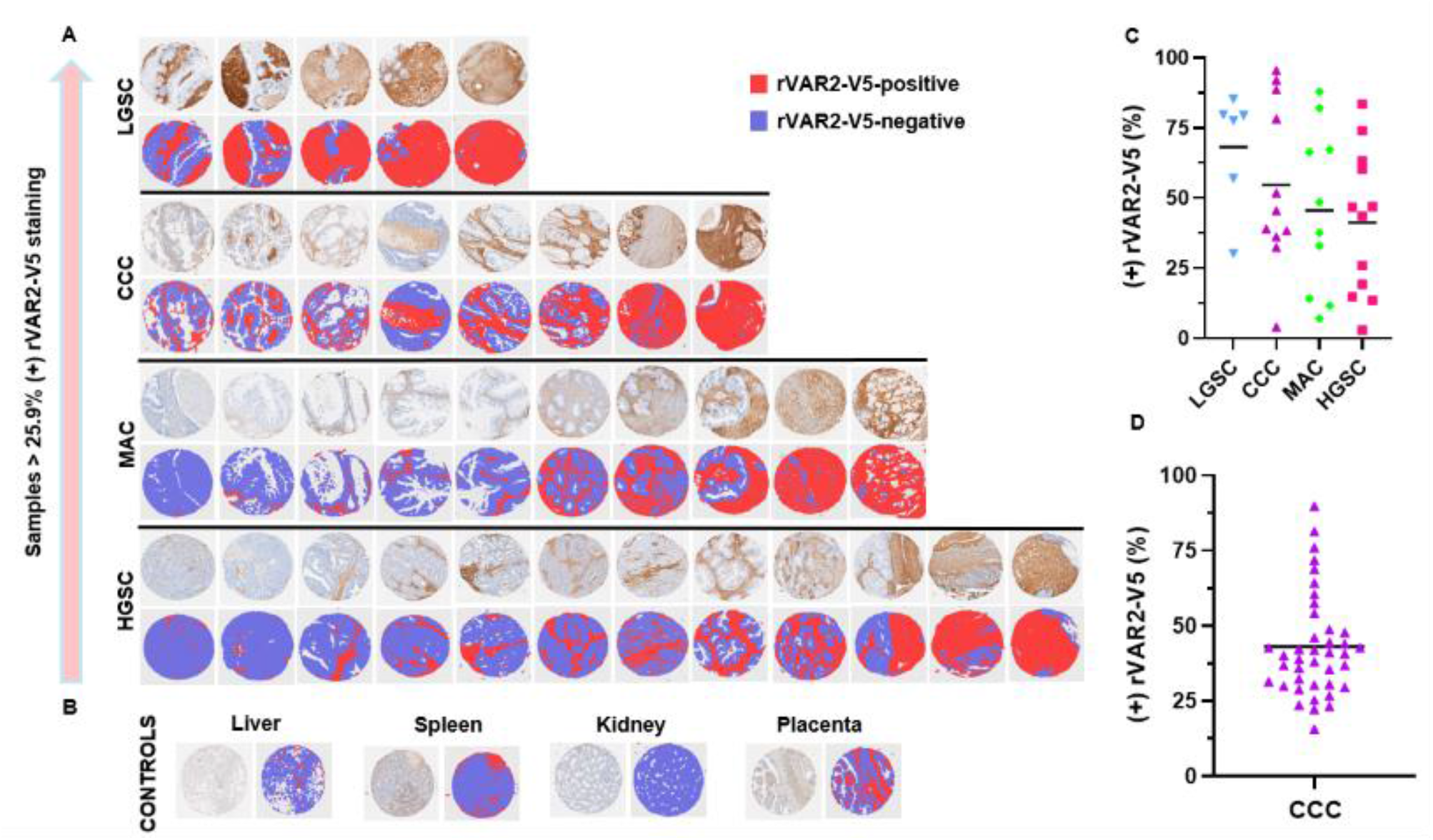
ofCS staining across ovarian cancer subtypes and normal tissues. (A) Representative ovarian cancer tissue microarray (TMA) cores from the UVA CHTN cohort, stained with rVAR2-V5, with corresponding digital image segmentations. Patient clinical histories were unavailable. (B) Normal liver, spleen, kidney, and placenta reference tissues stained with rVAR2-V5. In the digital segmentations in A and B, red indicates rVAR2-V5-positive tissue and blue indicates rVAR2-V5-negative tissue. (C) Percentage of tissue area positive for rVAR2-V5 in UVA CHTN ovarian cancer TMA samples grouped by histotypes. Each value represents the mean of four cores per patient sample. LGSC, low-grade serous carcinoma (*n* = 6); CCC, clear cell carcinoma (*n* = 11); MAC, mucinous adenocarcinoma (*n* = 10); HGSC, high-grade serous carcinoma (*n* = 12) (D) Percentage of tissue area positive for rVAR2-V5 in independent UPenn clear cell carcinoma TMA samples (n=40). Horizontal black lines in (C) and (D) indicate group means.

### Subhead 4: Triplatin is active in carboplatin-resistant models

Having identified a C4S-rich state associated with carboplatin resistance, we compared carboplatin with Triplatin, a cationic polynuclear platinum compound that interacts with sulfated GAGs (Fig. 5A). Under short-exposure treatment conditions, Triplatin showed greater potency than carboplatin in the ovarian clear cell carcinoma models OVTOKO and JHOC-5 and in ES2 cells (Fig. 5B-D).

**Fig. 5.**
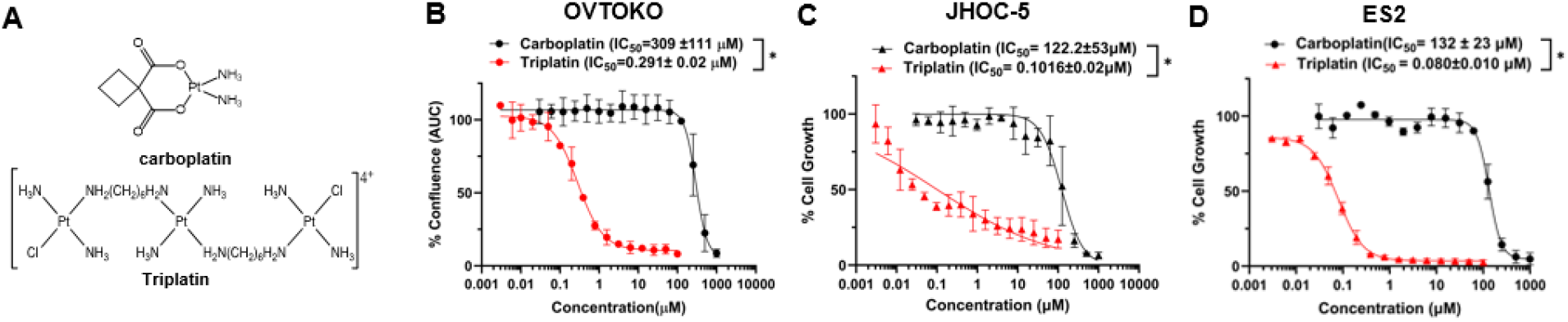
Triplatin is active in platinum-resistant ovarian cell lines. **(A)** Carboplatin and Triplatin chemical structures. **(B-D)** Cell growth inhibitory activity of carboplatin and Triplatin on human OVTOKO, JHOC5, and ES2 cancer cell lines; 1h treatment. ^*^p<0.05; Student’s t-test

### Subhead 5: GAG depletion shifts platinum pharmacology

To test whether GAG abundance directly modifies platinum activity, we generated GAG-deficient ES2 ovarian cancer cells by CRISPR-mediated knockout (KO) of *XYLT1* and *XYLT2* (XYLT), which encode xylosyltransferases that initiate GAG chain attachment to proteoglycan core proteins. We assessed C4S levels using rVAR2-V5. XYLT KO and chondroitinase ABC treatment each markedly reduced rVAR2-V5 binding relative to empty Cas9 vector (CTRL) cells, consistent with loss of intact C4S/ofCS-bearing GAG chains (Fig. 6A). Phase-contrast imaging showed that CTRL cells exhibited a more elongated, spindle-shaped morphology associated with a migratory phenotype, whereas XYLT KO cells were more compact and exhibited reduced cell spreading. Cell counts at 24 and 48 hours did not differ significantly between groups, indicating that the morphological differences were not accompanied by changes in short-term proliferation(Fig. 6B,C).

**Fig. 6.**
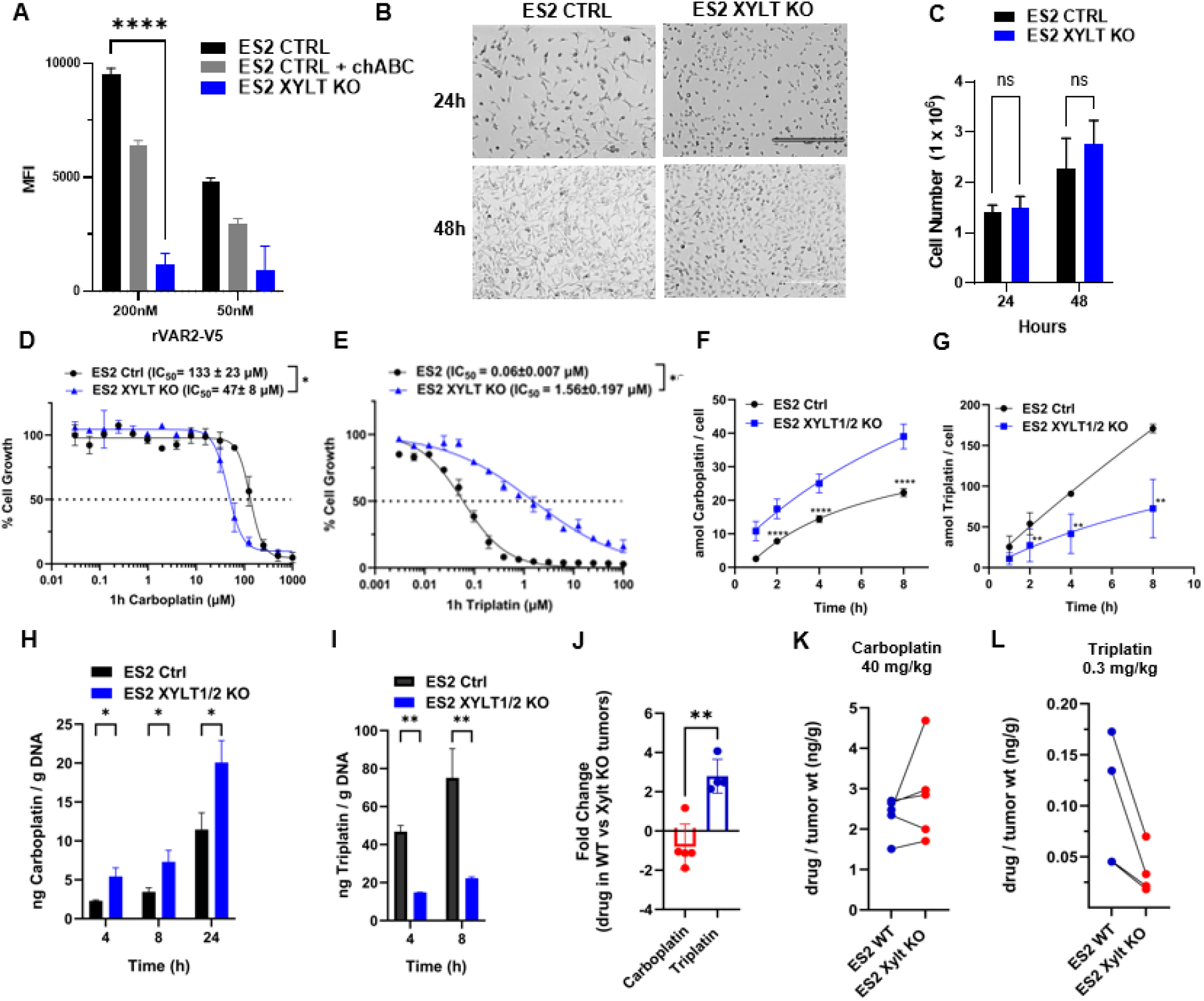
Changes in GAG levels inversely shift carboplatin and Triplatin sensitivity. **(A)** Flow cytometry of 200 nM and 50 nM rVAR2-V5 binding to ES2 CTRL cells, ES2 CTRL cells + chABC treatment, or XYLT KO cells. **(B)** Phase-contrast microscopy images (10× objective) of ES2-luc CTRL and XYLT KO cells captured at 24- and 48-hours post-seeding. **(C)** Cell counting; 1 x 10^6^ cells were seeded in 100mm dishes and counted at 24 and 48 h after seeding using a hemocytometer. **(D and E)** Cytotoxicity of carboplatin and Triplatin in ES2 CTRL and XYLT KO cell lines; 1h treatment. **(F and G)** Platinum cellular accumulation in ES2 CTRL and XYLT KO cells treated with 10 µM carboplatin or Triplatin for 1, 2, 4, and 8h. Platinum content was measured by inductively coupled plasma mass spectroscopy (ICP-MS) and normalized by number of cells. **(H and I)** Platinum-DNA adducts in ES2 CTRL and XYLT KO cells treated with 10 µM carboplatin or Triplatin 4, and 8h. **(J and K)** ES2-luc CTRL and XYLT KO tumors were implanted on the left and right flanks mice. Tumors were harvested after 24h treatment with 40 mg/kg i.p Carboplatin or 0.3 mg/kg i.p. Triplatin. Tumors were digested in nitric acid and platinum measured by ICP-MS. Statistical tests: Two-way ANOVA (A, C, F-J), Student’s t-test (D, E); ns = not significant, ^*^p<0.05, ^**^p<0.01, ^***^p<0.005, ^****^p<0.001

Functionally, GAG depletion sensitized ES2 cells to carboplatin but reduced sensitivity to Triplatin (Fig. 6D,E). This reciprocal drug-response pattern was mirrored pharmacologically: XYLT KO increased intracellular carboplatin accumulation and carboplatin-DNA adduct formation, whereas Triplatin accumulation and Triplatin-DNA adduct formation decreased (Fig. 6F-I). Matched flank tumor uptake experiments reproduced these directional effects *in vivo*, with carboplatin accumulating preferentially in GAG-deficient tumors and Triplatin accumulating preferentially in GAG-intact tumors (Fig. 6J-L). Parallel analyses in wt (CHO-K1) and GAG-deficient cells (pgsA-745) chinese hamster ovarian cell lines confirmed that the pharmacologic shift was not unique to ES2 cells (fig. S5)*(31)*. In an ES2-luc treatment survival experiment, Triplatin prolonged survival relative to carboplatin (fig. S6). Together, these data support a model in which a C4S-rich extracellular environment limits carboplatin accumulation but favors retention and activity of the GAG-targeting agent Triplatin.

### Subhead 7: GAG depletion suppresses tumor growth and dissemination

To evaluate whether GAG depletion alters ovarian tumor behavior in vivo, we monitored ES2 CTRL and XYLT KO subcutaneous xenografts by caliper measurement. XYLT KO tumors showed marked growth inhibition relative to CTRL tumors (Fig. 7A). Consistent with this growth phenotype, XYLT KO significantly prolonged endpoint-free survival (Fig. 7B). We next tested whether GAG depletion affected invasive behavior of tumor-derived cells in a 3D matrix model. Tumor cells isolated from CTRL and XYLT KO xenografts were dissociated, re-formed as multicellular spheroids, and embedded in collagen: fibrinogen hydrogel. CTRL spheroids displayed a more invasive, protrusive morphology, whereas XYLT KO spheroids remained more compact (Fig. 7C). Quantification confirmed that XYLT KO spheroids had significantly reduced perimeter and invasion index relative to CTRL spheroids, indicating decreased 3D migratory invasion after GAG depletion (Fig. 7D,E).

**Fig. 7.**
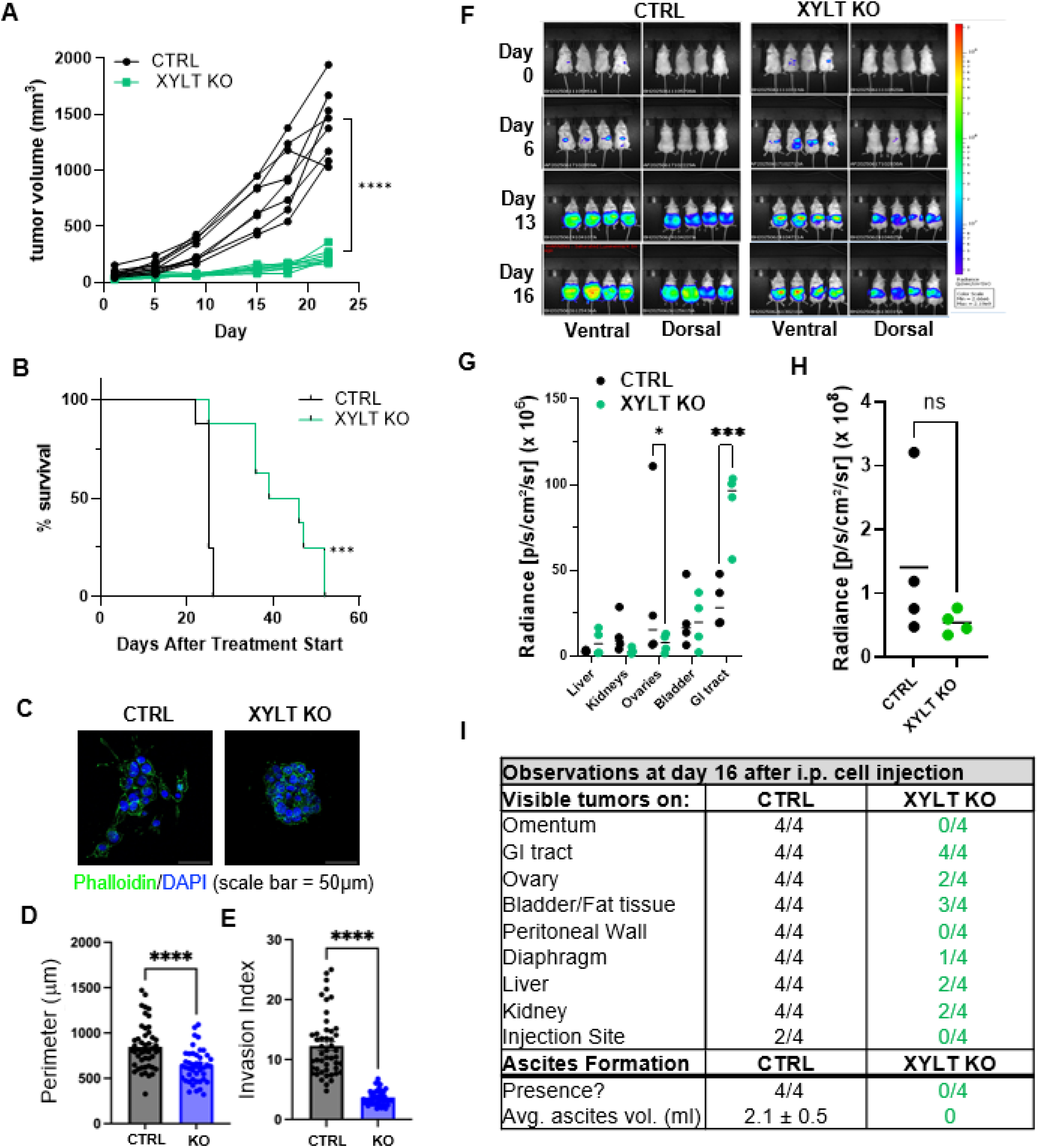
XYLT knockout suppresses tumor growth, prolongs endpoint-free survival, reduces spheroid invasion, and alters intraperitoneal dissemination and ascites formation in vivo. **(A,B)** NSG mice were subcutaneously injected with 1 × 10^6^ ES2 CTRL or XYLT KO cells, with n = 10 mice per group. **(A)** Tumor growth measured by caliper. ^****^p< 0.0001; two-way ANOVA **(B)** Kaplan–Meier curves showing the proportion of mice remaining below the prespecified tumor-volume endpoint of 2000 mm^3^. Mice were euthanized upon reaching this endpoint. ^***^p < 0.001; Log-rank **(C)** Representative confocal images of spheroids derived from CTRL and XYLT KO tumors and embedded in collagen: fibrinogen hydrogel. **(D**,**E)** Quantification of spheroid morphology and invasion, shown as **(D)** perimeter and **(E)** invasion index. n = 43– 48 spheroids per group, derived from 4–5 mice.^****^p < 0.0001; Welch’s t-test. **(F)** NSG mice were intraperitoneally injected with 1 × 10^6^ ES2 CTRL or XYLT KO cells, with n = 4 mice per group. Bioluminescent imaging was used to monitor tumor burden and distribution on days 0, 6, 13, and 16 after injection. Ventral and dorsal images are shown. Luciferase signal was acquired using the IVIS imaging system and quantified with Living Image software under consistent acquisition settings. **(G)** Ex vivo bioluminescent imaging of organs harvested on day 16. Total radiance was quantified for liver, kidneys, ovaries, bladder, and gastrointestinal tract using Living Image software.^*^p < 0.05, ^***^p < 0.001; two-way ANOVA **(H)** Whole-animal tumor-associated bioluminescent radiance, calculated for each mouse by summing ventral and dorsal radiance values and subtracting background signal (n = 4 mice per group). Each point represents an individual mouse. ns, not significant; Student’s t-test; **(I)** Gross necropsy summary of visible tumor distribution and ascites formation on day 16 after intraperitoneal injection. Organs and peritoneal sites were visually inspected for tumor presence at dissection, and ascites volume was measured.

To assess intraperitoneal dissemination, CTRL or XYLT KO cells were injected into abdominal cavity and monitored by serial bioluminescent imaging. Whole-animal imaging showed that XYLT KO altered tumor burden and distribution over time relative to CTRL cells (Fig. 7F). Ex vivo imaging on day 16 further demonstrated altered organ-associated tumor signal, with significant differences in ovarian and gastrointestinal tract radiance between CTRL and XYLT KO tumors (Fig. 7G). Whole-animal radiance followed a similar trend to the subcutaneous tumor-growth findings, with lower mean values in XYLT KO mice, although the difference was not statistically significant (Fig. 7H). Gross necropsy confirmed that XYLT KO reduced the frequency of visible tumor involvement across multiple peritoneal sites and eliminated detectable ascites formation, whereas all CTRL-bearing mice developed ascites and associated weight gain (Fig. 7I, fig. S7). Together, these findings indicate that the GAG-rich phenotype is not merely a passive biomarker, but is linked to ovarian tumor growth, invasive behavior, intraperitoneal distribution, and ascites formation.

### Subhead 8: Tissue C4S/ofCS staining marks Triplatin-sensitive PDX

Finally, to evaluate rVAR2-V5 staining as a pathology-compatible histologic readout of the C4S-rich glycomics phenotype, we compared it with direct GRIL-LC/MS measurements in ovarian cancer PDX tumors. Whole-slide digital pathology showed that rVAR2-V5 positivity tracked with the C4S-rich state identified by GRIL-LC/MS, with the highest staining in carboplatin-resistant models and the lowest staining in the carboplatin-sensitive model. (Fig. 8A-C). When these same models were treated *in vivo*, high-C4S/high-rVAR2 tumors were selectively sensitive to Triplatin, whereas low-C4S tumors were relatively more sensitive to carboplatin (Fig. 8D). Orthogonal staining with a C4S-directed antibody *(24)* yielded concordant spatial patterns and relative ranking across models (fig. S8), and the structurally related compound BBR3571 *(38)* showed a response pattern similar to Triplatin (fig. S9). These findings indicate that tissue-level detection of the glycan state can recapitulate the translationally relevant biology identified by direct glycomics and can distinguish tumors likely to respond preferentially to Triplatin rather than carboplatin.

**Fig. 8.**
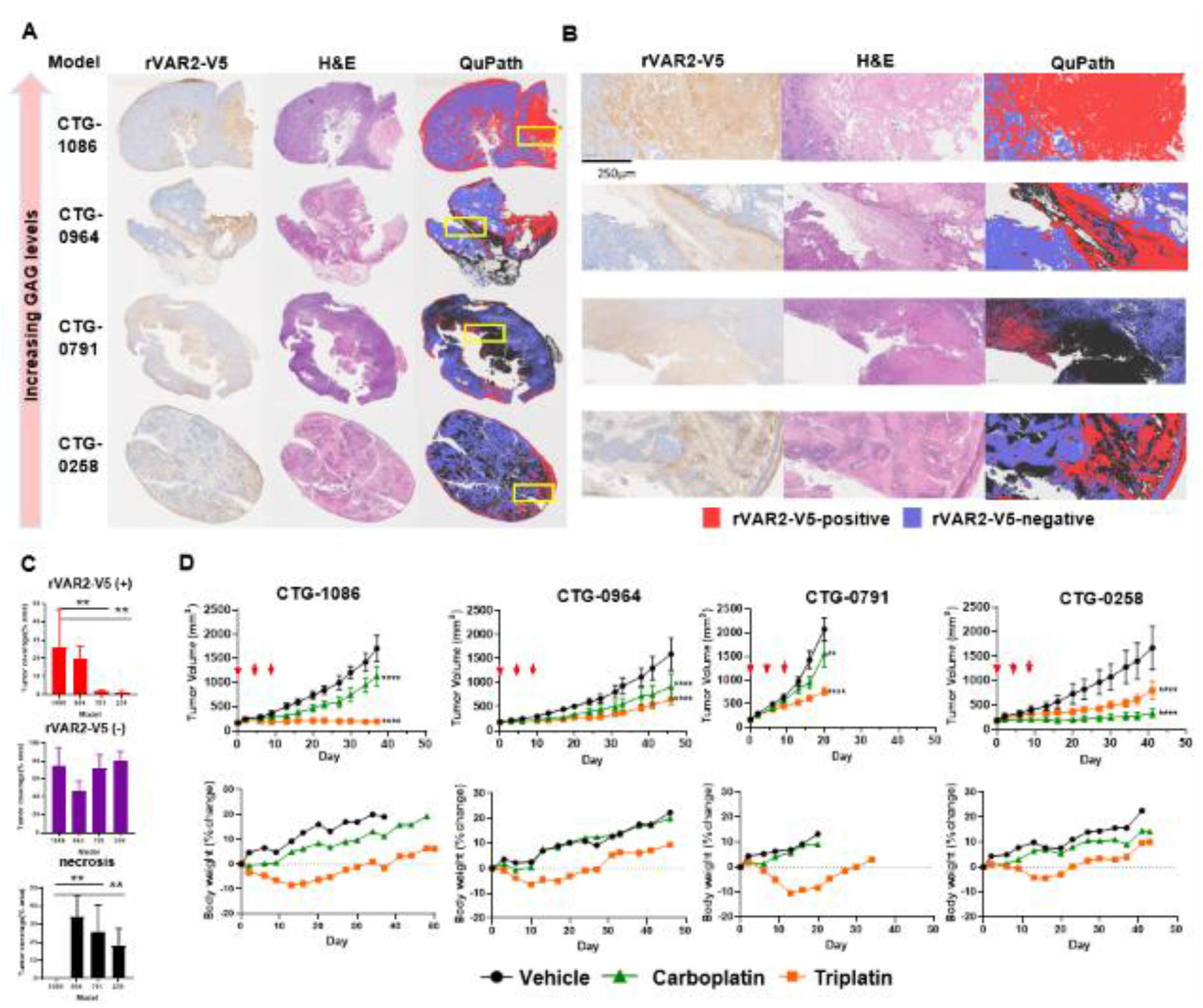
Triplatin inhibits tumor growth in carboplatin-resistant PDX expressing high levels of C4S. **(A and B)** Representative images of rVAR2-V5 and H&E OC PDX staining and Qupath analysis. **(C)** Qupath segmentation of rVAR2-V5 (+), rVAR2-V5 (-), and necrotic tumor area in OC PDX models. **(D)** Sensitivity of OC PDX models to Triplatin and carboplatin. OC PDX models were treated i.p. with carboplatin (40 mg/kg) or Triplatin (0.3 mg/kg) on days 0, 4 and 8 (orange arrows). ^**^p<0.01, ^****^p<0.0001, 2-way ANOVA, Tukey’s test.

## DISCUSSION

Our findings identify a stromal C4S/ofCS-rich extracellular state as a functionally important determinant of platinum response in ovarian cancer. The central advance is not simply the association of one biomarker with one drug, but the demonstration that the biologically relevant determinant resides in a matrix phenotype that is poorly captured by conventional transcriptomic inference. Across ovarian PDX models, direct glycomics profiling revealed that carboplatin resistance is associated with enrichment of C4S, whereas tissue-level rVAR2-V5 staining provides a clinically tractable readout of this same state. In parallel, functional perturbation of GAG biosynthesis shifted platinum accumulation and DNA platination in opposite directions for carboplatin and Triplatin, establishing that the glycan phenotype is mechanistically linked to drug disposition rather than being a passive correlate.

A major implication of this work is methodological. Transcriptomics can localize proteoglycan-producing cell states and, in our analyses, nominate stromal fibroblasts as a dominant source of candidate scaffolds such as VCAN and BGN. However, transcript abundance alone cannot define the abundance, length, or sulfation pattern of the mature GAG chains, because GAG biosynthesis is non-template-driven and determined by the coordinated action of multiple enzymes. In this setting, the active biology resides in the mature glycan structure, not in any single transcript. This explains why TCGA-style transcriptomic anchoring is insufficient for clinical translation of this biomarker and why direct glycomics or tissue-level biomarker assays are required to capture the extracellular phenotype that shapes platinum response.

The stromal localization of the biomarker-positive state is also biologically informative. Our tissue studies, single-cell analyses, and XYLT perturbation models collectively support a framework in which fibroblast-associated proteoglycan programs generate a matrix environment that affects both malignant phenotypes and intratumoral drug distribution. XYLT KO altered intraperitoneal spread and decreased ascites formation *in vivo*, consistent with broader roles for GAG-rich stroma in ovarian tumor progression and dissemination. Within this mechanistic model, a C4S/ofCS-rich microenvironment may reduce effective carboplatin penetration or local availability while favoring retention and activity of the GAG-binding agent Triplatin. Although the precise chemical and biophysical basis of these effects remains to be fully defined, the directional pharmacologic consequences were reproducible across cellular, xenograft, and PDX systems.

These findings have translational consequences for patient selection. High C4S/ofCS expression was common in ovarian cancer subtypes that are intrinsically less responsive to platinum, including clear cell and low-grade serous carcinomas, and tissue-level detection identified PDX tumors that were selectively sensitive to Triplatin. This argues for a biomarker-guided development strategy in which direct tissue measurement of the glycan state is used to identify patients unlikely to derive durable benefit from carboplatin and more likely to benefit from Triplatin or related GAG-targeting platinum agents. Because the biomarker-positive state appears to be rooted in stromal biology, this strategy could be particularly valuable in tumors where extracellular matrix composition constrains drug distribution.

Several limitations remain. First, larger clinically annotated cohorts will be needed to refine and prospectively validate a clinically actionable cut point for tissue staining. Second, although the transcriptomic analyses strongly support fibroblasts as a major cellular source of proteoglycan core-protein transcripts, they do not directly measure the abundance or sulfation of the attached GAG chains. Integrating spatial transcriptomics, glycomics, and histopathology in matched human tumors could determine whether fibroblast expression of these genes correlates with C4S abundance and its localization within the tumor stroma. Third, the conventional model of platinum drug activation emphasizes ligand substitution, including the formation of aqua species through substitution of leaving group (Cl-, CBDCA or oxalate in the case of oxaliplatin) within the intracellular environment, facilitating DNA crosslinking and cytotoxicity. However, our findings suggest that extracellular matrix components, particularly GAGs, may function as an overlooked “sink” of oxygen donors for platinum species, altering their local bioavailability and reactivity*(39)*. While our data demonstrates functional effects on platinum disposition, additional work will be required to dissect the relative contributions of extracellular sequestration, altered penetration, transport processes, and microenvironment-dependent drug activation. Even with these limitations, the current results establish an important conceptual advance: the extracellular glycan state is a non-genetic determinant of platinum response that can be measured directly and targeted therapeutically.

The findings presented here support the development of C4S/ofCS-directed treatment strategies in ovarian cancer, particularly for patients with recurrent platinum-resistant disease or those with inherently resistant histologic subtypes*(40)*. Translation into clinical practice will first require prospective studies evaluating C4S/ofCS staining in biopsy specimens to determine whether expression reliably predicts Triplatin sensitivity and can support patient stratification. Triplatin’s unique GAG-binding properties suggest a broader therapeutic application beyond ovarian cancer, particularly in other GAG-rich tumors such as pancreatic cancer, triple-negative breast cancer and bladder cancer, where similar resistance patterns have been observed*(15, 19, 20, 24)*. If C4S/ofCS functions as a barrier to carboplatin entry, targeting C4S/ofCS, enzymatically or with competitive inhibitors, may enhance carboplatin efficacy. Another approach that ovarian cancer patients may benefit from is the use of C4S/ofCS-targeting modalities-such as antibody-drug conjugate or bispecific antibodies, which have demonstrated safe and potent anti-tumor activity in multiple animal models across diverse cancer types *(19, 24, 41, 42)*. Future studies should investigate whether combining carboplatin with GAG-modulating agents can further optimize treatment outcomes. Given the growing emphasis on biomarker-driven treatment selection in oncology, incorporating C4S/ofCS profiling into precision medicine strategies may allow for more personalized and effective treatment regimens, particularly in platinum-resistant ovarian cancer. Future clinical studies will be essential to refining C4S/ofCS-based patient stratification, optimizing treatment selection, and developing next-generation platinum compounds that exploit tumor-specific GAG expression to overcome resistance.

## MATERIALS AND METHODS

### Study Design

The objective was to determine whether glycosaminoglycans (GAGs), particularly chondroitin-4-sulfate/oncofetal chondroitin sulfate structures, define a functionally relevant extracellular state that modulates platinum response in ovarian cancer and can serve as a translational biomarker for Triplatin sensitivity. We combined quantitative glycomics of PDX tumors, public single-cell transcriptomic analysis to nominate source cell populations, *in vitro* and *in vivo* perturbation of GAG biosynthesis, platinum uptake and DNA adduct measurements, xenograft and PDX efficacy studies, and tissue-based biomarker profiling by immunohistochemistry and digital pathology.

### Reagents

Triplatin was synthesized according to published procedures *(43)*. The malarial protein rVAR2 tagged with a V5 peptide (rVAR2-V5) was produced as described previously*(19)* .Carboplatin (cat# PHR3417) and chondroitinase ABC were purchased from Sigma. Anti-V5 antibody (cat. # 96025) and anti-V5-Alexa Fluor 647 conjugate antibody (cat. # 451098) were purchased from ThermoFisher.

### Glycan profiling (GRIL-MS; Glycan Reductive Isotope Labeling Mass Spectrometry)

GAG isolation, purification, and aniline tagging were performed as previously described *(44)*, with minor modifications. Flash-frozen PDX tumor tissues were pulverized and homogenized in protease digestion buffer. Samples were incubated overnight at 37 °C with protease and centrifuged at 14,000 rpm for 20 minutes. The resulting supernatant containing released GAGs was applied to a DEAE ion exchange column, and bound GAGs were eluted with 2 M NaCl. Eluted GAGs were desalted using PD-10 Sephadex G-25 desalting columns (GE Healthcare), lyophilized, and resuspended in digestion buffer specific for chondroitin sulfate (CS) or heparan sulfate (HS). CS-containing samples were digested with a cocktail of chondroitinase ABC, while HS-containing samples were digested with a mixture of heparinase I, II, and III (IBEX). Digestions were performed at 37 °C for 18 hours. Following digestion, samples were passed through 10,000 molecular weight cutoff filters to remove enzymes and undigested high-molecular-weight GAG chains. The filtrate containing disaccharide products was collected and dried under vacuum. The dried disaccharides were dissolved in aniline and derivatized by reductive amination using freshly prepared sodium cyanoborohydride in a dimethyl sulfoxide:acetic acid (7:3, v/v) mixture. Labeling reactions were incubated at 65 °C for 1 hour, followed by 16 hours at 37 °C. Samples were then dried in a speed vacuum concentrator at room temperature and submitted to the UCSD GlycoAnalytics Core for glycan analysis. Aniline-tagged disaccharides are separated on a C18 column using an ion pairing solvent mixture and analyzed by linear trap quadrupole mass spectrometry*(32)*.

### Single-cell transcriptomic analysis

Public single-cell RNA-seq datasets were used to nominate cellular sources of proteoglycan core proteins associated with the C4S-rich phenotype. Analysis included normal ovarian reference data from the Tabula Sapiens consortium and publicly available ovarian or fallopian tube tumor datasets accessed through CELLxGENE Discover, including matched primary and metastatic sites when available*(35–37)*. Cell-type annotations supplied with the datasets were used to compare expression of proteoglycan genes across epithelial, stromal, endothelial, and immune compartments. These analyses were used to identify candidate source cell populations for matrix-associated proteoglycan programs.

### Cell lines and culture conditions

CHO-K1 and CHO pgsA-745 cells were purchased from ATCC and cultured in DMEM/F-12 medium supplemented with 10% fetal bovine serum (FBS) and 1% penicillin/streptomycin. OVTOKO-luc, JHOC-5-luc, and ES2-luc cells were provided by Dr. Ronny Drapkin (University of Pennsylvania). ES2-luc cells were cultured in McCoy’s 5A medium (Invitrogen) supplemented with 10% FBS and 1% penicillin/streptomycin (Gibco). All cells were grown at 37 C in a humidified atmosphere with 5% CO2 and were routinely tested for mycoplasma contamination by PCR.

### CRISPR/Cas9 XYLT1/2 KO cell line generation

GAG-deficient ES2-luc cell lines were generated according to published procedures *(45)*. Parental ES2-luc cells were transfected with either Cas9 plasmid (KO) or empty DNA backbone (CTRL) plus *XYLT1* and *XYLT2* gRNA plasmids. Following transfection, GFP-positive cells were single-cell sorted into 96-well plates and expanded for 4 weeks. CTRL and KO cells were then screened by flow cytometry using rVAR2-V5 to identify GAG-deficient clones. Genomic alterations in KO cells were confirmed by PCR amplification of XYLT1 and XYLT2, cloning into TOPO TA sequencing plasmids, and Sanger sequencing (Eurofins) with the M13F primer. Primers were XYLT1 forward, gtttctctcccctcttctccag; XYLT1 reverse, atgtcacacttagggggctg; XYLT2 forward, cccagtgatgtgtctgcatc; and XYLT2 reverse, gtatctccgtgtggtgggaag. Each cell line was sequenced four times for each gene.

### Cellular viability (MTT assay)

Cells were seeded in 96-well plates at 1 × 10^3^ cells/well in 100 µL of complete growth medium and incubated for 24 h at 37 °C in an incubator with 5% CO_2_. Cells were treated with drug or vehicle for 1h, washed once with PBS, and returned to fresh growth medium. When control wells reached confluence, medium was replaced with 100 µL of MTT reagent (0.5 mg/ml in growth medium), and plates were incubated for 3 h at 37 °C. MTT solution was removed, formazan crystals were solubilized in 100 µL of DMSO, and plates were incubated for 1 h at room temperature. Absorbance was measured at 570 nm using a BioTek Synergy microplate reader, with background subtraction. Viability was calculated relative to untreated control wells, set to 100%. Data were analyzed in GraphPad Prism and are reported as mean +/-SD from at least two independent experiments with four technical replicates each.

### C4S/ofCS quantification (Flow cytometry)

Cells were harvested using Versene and washed twice in cold PBS. Cells (1.5 x 10^6) resuspended in PBS containing 0.5% BSA and kept on ice. Cells were incubated with 50 or 200 nM rVAR2-V5 for 30 min at 4°C, washed twice with PBS, and incubated with anti-V5-Alexa Fluor 647 antibody (1:400; Invitrogen, cat. no. 451098). Binding was analyzed using a BD Fortessa flow cytometer. For enzymatic GAG removal, ES2 control cells were washed in PBS and incubated with 0.1 U/ml chondroitinase ABC in buffer containing 50 mM Tris-HCl (pH 8.0), 60 mM sodium acetate, and 0.02% BSA for 30 min at room temperature before washing and rVAR2-V5 staining.

### Cellular platinum levels *in vitro* (ICP-MS)

Intracellular platinum content was determined as previously described *(29)*, with minor modifications. Cells were seeded onto 100-mm tissue culture dishes and allowed to adhere for 24 h at 37 °C in an incubator with 5% CO_2_. At each timepoint, cells were harvested by trypsinization, washed with PBS to remove extracellular drug, and counted. Cell pellets were digested in trace-metal–grade nitric acid at room temperature for 72 hours. The digested samples were filtered through 0.22 µm PVDF filters and analyzed for platinum content using ICP-MS. Platinum concentrations were quantified against a standard curve generated using platinum standards of known concentration and normalized to total cell number. Data are presented as intracellular platinum per 10^6^ cells. Cell pellets were digested in trace-metal-grade nitric acid for 72 hours at room temperature. Digests were filtered through 0.22-micrometer PVDF filters and analyzed for platinum by ICP-MS against platinum standards. Platinum content was normalized to total cell number and is presented as platinum per 10^6 cells.

### Cellular platinum-DNA adducts *in vitro* (ICP-MS)

Cells were seeded onto 100 mm dishes, allowed to attach for 24 h, and treated with 10 μM carboplatin or Triplatin. At the indicated times, cells were harvested by trypsinization and washed repeatedly with PBS. DNA was extracted using the Qiagen DNeasy Blood and Tissue kit and digested in trace-metal-grade nitric acid for 72 hours at room temperature. Digests were filtered through 0.22-micrometer PVDF filters and analyzed for platinum by ICP-MS against platinum standards. Platinum content was normalized to input DNA mass and is presented as platinum per gram DNA.

### Tumor platinum levels in vivo (ICP-MS)

For in vivo platinum uptake studies, ES2-luc CTRL/XYLT KO tumors or CHO-K1/CHO pgsA-745 tumors were excised 24 hours after a single i.p. dose of carboplatin (40 mg/kg) or Triplatin (0.3 mg/kg). Tumors were weighed, minced, and digested in trace-metal-grade nitric acid (2 ml per 100 mg tissue) for 72 hours at room temperature. Digests were clarified through 0.22-micrometer PVDF filters and analyzed by ICP-MS against platinum standards. Data are reported as micrograms or nanograms platinum per gram wet tumor.

### PDX models and in vivo drug efficacy

Ovarian cancer PDX tumors with prior carboplatin sensitivity annotation were obtained through Champions Oncology for glycan profiling and efficacy studies. Drug efficacy studies were performed by Champions Oncology in four ovarian cancer PDX models (CTG-1086, CTG-0964, CTG-0791, and CTG-0258). PDX fragments of approximately 3 mm^3^ were implanted subcutaneously into the flank of 8-to 10-week-old NSG mice. Mice were randomized when tumor volume reached 100 to 200 mm3. Carboplatin (40 mg/kg i.p.), Triplatin (0.3 mg/kg i.p.), BBR3571 (0.3 mg/kg i.p.) or saline was administered on days 0, 4, and 8 (q4d x 3). Tumors were measured two to three times per week by caliper, and volume was calculated as (L x W^2)/2. Percent tumor growth inhibition (TGI) was calculated at the indicated study day as 100 x [1 - (DeltaT/DeltaC)], where DeltaT and DeltaC mean tumor volume changes from baseline in treated and control cohorts. Growth curves are shown as mean +/-SEM. BBR3571 was evaluated as an additional treatment arm in the supplemental PDX study shown in fig. S9 using the same treatment schedule.

### ES2-luc xenograft studies

Mouse experiments were conducted by VCU’s Cancer Mouse Models Core and approved by the VCU Institutional Animal Care and Use Committee under the Animal Protocol AD10002979, in compliance with VCU institutional guidelines for the care and use of animals. For flank drug-uptake studies, ES2-luc CTRL and XYLT KO cells (5 x 10^6 in 1:1 PBS: Matrigel; 100 microliters) were injected into contralateral flanks of female NSG mice. When tumors reached approximately 200 mm3, mice received one dose of carboplatin (40 mg/kg) or Triplatin (0.3 mg/kg) i.p.; tumors were harvested 24 h later for ICP-MS. For flank tumor-growth studies, ES2-luc CTRL or XYLT KO cells (1 x 10^6) were injected subcutaneously into female NSG mice (n = 10 per group), tumor volume was measured by caliper, and mice were euthanized when tumor volume reached 2000 mm3. For the ES2-luc treatment survival experiment, mice received carboplatin (40 mg/kg i.p.) or Triplatin (0.3 mg/kg i.p.) on days 7, 11, and 15. For the carcinomatosis model, ES2-luc CTRL or XYLT KO cells (1 x 10^6) were injected intraperitoneally into female NSG mice (n = 4 per group), and disease progression was monitored by bioluminescent imaging, body weight, gross tumor distribution, and ascites volume.

### Tumor-derived spheroid invasion assay

Tumor cells were isolated from ES2-luc CTRL and XYLT KO xenograft tumors. Tumors were dissociated to single-cell suspensions, and cells were plated in low-adhesion wells at 150,000 cells/mL in culture medium containing 2% Matrigel (Corning, CB-40230) to generate multicellular spheroids for 24 h. Spheroids were collected, resuspended in type I rat-tail collagen: fibrinogen hydrogel (2 mg/mL collagen and 5 mg/mL fibrinogen), and cultured in 8-well chamber slides (Cellvis, C8-1-N). Spheroids were incubated for 18–24 h to allow invasion and then fixed in 4% paraformaldehyde. Immunofluorescence staining and 3D confocal imaging were performed on a Zeiss LSM 980 microscope (40× water objective, 0.6× zoom; Nanomaterials Core Characterization Facility at VCU) to visualize F-actin with Alexa Fluor phalloidin (Invitrogen), nuclei with DAPI (Invitrogen, D1306), and extracellular matrix fibers by confocal reflectance. Actin images for each spheroid were compiled into z-projections and thresholded in FIJI to generate binary masks. Area and perimeter were measured in FIJI, and invasion index was calculated as perimeter^2^/(4π × area), as described previously *(41)*.

### Tissue microarray cohorts

OvCa2 multi-histology TMA samples were obtained from the University of Virginia Cooperative Human Tissue Network (0.6-mm cores, quadruplicate cores per case). An independent clear cell carcinoma TMA was provided by the Penn Medicine BioTrust. Clinical histories were not available for outcome analysis.

### Immunohistochemistry

IHC staining for C4S/ofCS using rVAR2-V5 was performed in the VCU Tissue and Data, Acquisition, and Analysis Core on a Leica Bond RX autostainer. Tissues were fixed in 10% neutral buffered formalin for at least 5 days before paraffin embedding and sectioning. Slides were baked at 60 C for 60 to 120 min, deparaffinized, and rehydrated through graded ethanols to distilled water. Antigen retrieval was not required for rVAR2-V5 staining. rVAR2-V5 was diluted in casein buffer to 500 pM, and 150 microliters was applied to each slide for 45 min at room temperature in a humidified chamber, followed by three washes. Bound peptide was detected with mouse anti-V5 antibody (R960-25, ThermoFisher; 1:700 in casein buffer, 45 min, room temperature), Leica post-primary rabbit anti-mouse antibody (8 min), and goat anti-rabbit HRP conjugate (8 min), with washes between steps. Slides were developed with 3,3-diaminobenzidine (DAB; Leica kit) for 10 min, counterstained with hematoxylin, dehydrated, cleared, and mounted. Positive controls included tumors with validated high and low C4S/ofCS expression, and negative controls omitted rVAR2-V5. Stained slides were scanned at 20x magnification on a Phenoimager HT whole-slide scanner (Quanterix) and analyzed in QuPath v0.7.0. Orthogonal C4S antibody staining with the C9 antibody was performed on serial PDX sections as described previously (22), and percent positive staining area was quantified in QuPath.

### Digital pathology and image analysis (Qupath)

The rVAR2-V5 proportion score (PS) was defined as the percentage of tumor area positive above a fixed DAB optical density threshold determined on control tissues and applied batch-wise. For each TMA case, PS was averaged across available cores. Artifacts were excluded manually. A conservative cut point of PS >= 26% was selected a priori from PDX-anchored correlation analyses linking rVAR2-V5 area fraction to Triplatin-versus-carboplatin response separation and was then applied to TMAs to estimate prevalence by subtype. For tumor-stroma analysis, QuPath pixel classifiers were trained to segment tumor epithelium and stroma on matched H&E and rVAR2-V5-stained sections. Tumor-stroma ratio (TSR) was calculated from segmented areas, and tumors were categorized as stromal-rich when TSR < 1.0.

## Statistical analysis

Statistical analyses were performed using GraphPad Prism 10. Assay-specific tests are indicated in the figure legends. One-way or two-way analysis of variance (ANOVA) was used for comparisons across multiple groups, with Tukey or Dunnett multiple-comparison tests as appropriate. Unpaired or paired t-tests were used for two-group comparisons as indicated. Kaplan-Meier survival curves were analyzed by log-rank test. Fisher’s exact test was used for categorical tumor-stroma comparisons. P values < 0.05 were considered statistically significant.

## List of Supplementary Materials

*Figs. S1 to S9*

## Acknowledgements

Services and products in support of the research project were generated by the VCU Massey Comprehensive Cancer Center Cancer Mouse Models Core, Tissue and Data, Acquisition and Analysis Core, and Flow Cytometry Core supported, in part, with funding from NIH-NCI Cancer Center Support Grant P30 CA016059.

We thank Biswa P. Choudhury and the UCSD GlycoAnalytics Core for assistance and support of the initial GRIL-MS analysis.

We thank the University of Pennsylvania Perelman School of Medicine Ovarian Cancer Research Center Tumor BioTrust Collection Core Facility, RRID:SCR_022387 for supplying the clear cell carcinoma TMA.

## Funding

Commonwealth Health Research Board grant

Virginia Innovation Partnership Corporation grant

Massey Comprehensive Cancer Center

BRIDGE Translational Excellence Program grant (NNF23SA0087869) for EEVC

Terry Fox Research Institute Program Project Grant (TFRI-PPG) (#GR026025) for MD

## Author contributions

Conceptualization: EJP, JEK, LL, NPF

Methodology: EJP, JBMT, JDH, JEK, RJW, TMC, EEV, AS,

Investigation: EJP, JDH, JEK, RJW, SPD, MGD, AB, EEV

Visualization: EJP, JEK, ARSB,

Funding acquisition: EJP, JEK, LL, NPF

Project administration: EJP, LL, NPF

Supervision: EJP, JEK, LL, NPF,

Writing – original draft: EJP

Writing – review & editing: EJP, JEK, LL, NPF

## Data and materials availability

All data are available in the main text or the supplementary materials.

