## Supplemental Figures for "Glycomics Profiling Reveals a Stromal Chondroitin-4-sulfate State that Shapes Platinum Response in Ovarian Cancer"

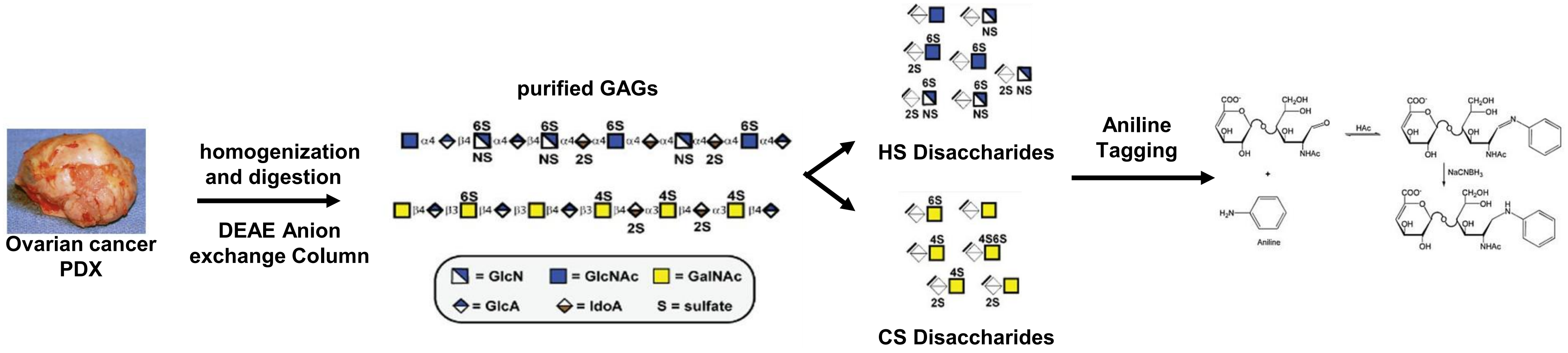


**Fig. S1 GRIL-MS disaccharide analysis of glycosaminoglycans**. Overview of the workflow for isolation, purification, and reductive isotopic labeling for MS analysis.

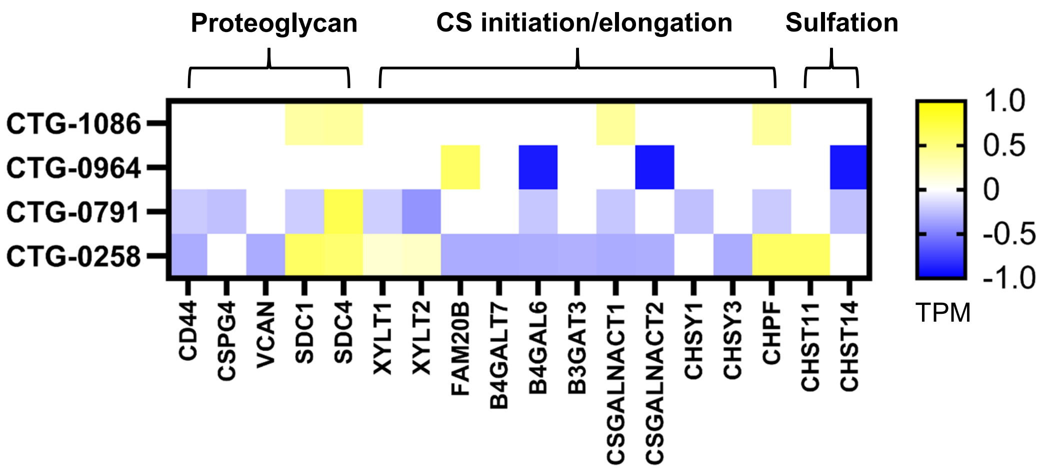


**Fig. S2. Expression of proteoglycan and chondroitin/dermatan sulfate biosynthesis genes in ovarian cancer PDX models.** Heatmap of selected genes in four ovarian cancer patient-derived xenograft (PDX) models: CTG-1086, CTG-0964, CTG-0791, and CTG-0258. Bulk RNA-sequencing data were provided by Champions Oncology. Rows represent PDX models, and columns represent genes grouped by their involvement in proteoglycan core protein expression, chondroitin sulfate/dermatan sulfate (CS/DS) chain initiation and elongation, or sulfation. Expression is displayed as $\log_2$-transformed transcripts per million (TPM), with yellow and blue indicating higher and lower values, respectively.

| Gene | High-expression *n* | Median PFI: high, months | Median PFI: others, months | Log-rank P |
|---|---|---|---|---|
| HSPG2 | 15 | 9.8 | 17.0 | 0.0012 |
| CSGALNACT2 | 14 | 12.7 | 17.0 | 0.0077 |
| VCAN | 12 | 12.3 | 16.1 | 0.0135 |
| CHST15 | 17 | 10.7 | 16.1 | 0.0282 |
| CHSY3 | 19 | 13.0 | 17.0 | 0.0446 |
| BCAN | 5 | 6.0 | 15.7 | 0.0489 |

**Fig. S3. CS-associated gene expression and progression-free interval in carboplatin-treated ovarian cancer.** RNA-seq expression data and treatment timelines were obtained from the Ovarian Serous Cystadenocarcinoma (TCGA, PanCancer Atlas) study in cBioPortal. Clinical annotations were supplemented with matched TCGA Firehose Legacy records, and event and censoring data were obtained from the TCGA Pan-Cancer Clinical Data Resource. The table summarizes genes for which high expression was associated with shorter progression-free interval (PFI) in 185 stage III–IV TCGA ovarian serous carcinoma patients with documented adjuvant carboplatin as their first recorded platinum therapy. High expression was defined as a diploid-reference mRNA z-score >1.5 and compared with $z \leq 1.5$. Columns show high-expression group size, Kaplan–Meier median PFI for each group, and two-sided log-rank P values. PFI was measured from first carboplatin administration to the TCGA-CDR event or censoring date; events included new tumor events and death with tumor. Displayed genes were selected from a 59-gene candidate panel using nominal $P < 0.05$ and an adverse association with high expression.

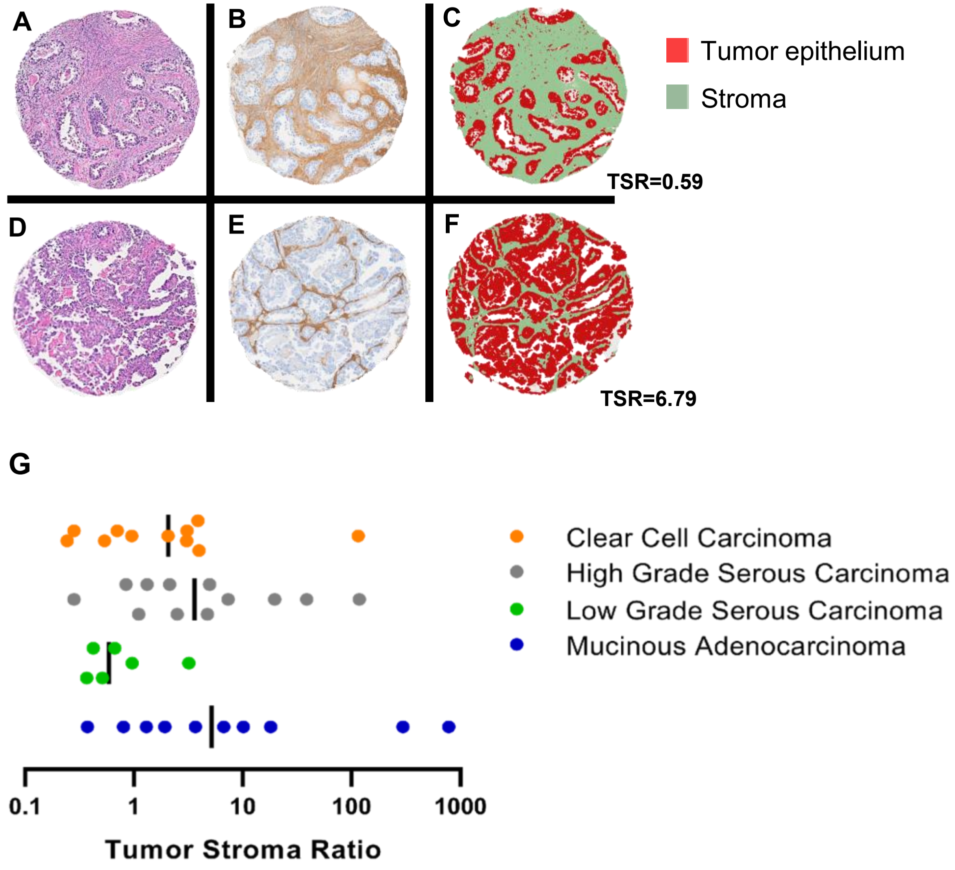


**Fig. S4. C4S staining colocalizes with tumor stroma.** Images display representative OC tumors with high stroma **(A-C)** and low stroma **(D-F)** comparing matched H&E staining **(A,D)**, rVAR2-V5 staining **(B,E)** and segmented images of tumor epithelium (red) and stroma (green) **(C,F)**. Using the pixel classifier algorithm in QuPath, segmented images were used to calculate the tumor stroma ratio (TSR). TMA shows 36% (14/39) with TSR<1.0 and 64% (25/39) with TSR > 1.0. **(G)** Distribution of TSR in various subtypes of ovarian cancer. Stromal-rich tumors (TSR<1.0); 45% CCC (5/11), 17% HGSC (2/12), 83% LGSC (5/6), 20% MAC (2/10). Tumor epithelium and stroma segmented using the pixel classifier algorithm in QuPath based on rVAR2 staining. TMA CHTN OvCa2 provided by UVA

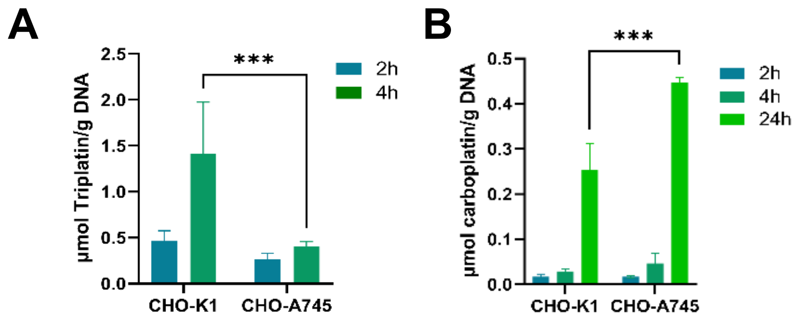


**Fig. S5 Effects of GAG levels on cellular DNA platination A, B.** Levels of Pt-DNA adducts in K1 and A745 cells treated with 10 µM Triplatin or 100 µM carboplatin. Platinum content in isolated DNA was measured by ICP-MS and normalized to DNA quantity. ***p <0.001, two-way ANOVA

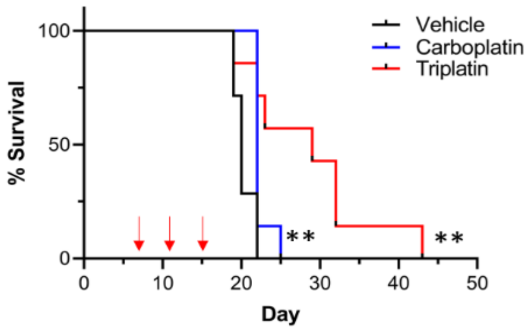


**Fig. S6 Increased survival in ES2-luc mouse model treated with Triplatin.** Survival analysis of ES-2-luc orthotopic model treated with carboplatin (40 mg/kg) and Triplatin (0.3 mg/kg) given i.p. on day 7, 11, and 15 (red arrows). **p<0.01, Log-rank

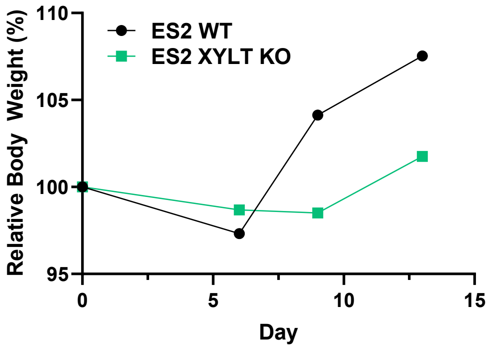


**Fig. S7 XYLT1/2 KO reduces weight gain caused by ascites formation *in vivo*.** Body weight measured on day 0, 6, 9 and 13. after i.p. injection with 1x$10^6$ ES2 WT or ES2 XYLT KO cells (n=4 mice per group). Relative Body Weight = mean(T/T0) * 100%. T0 = initial weight; T = current weight.

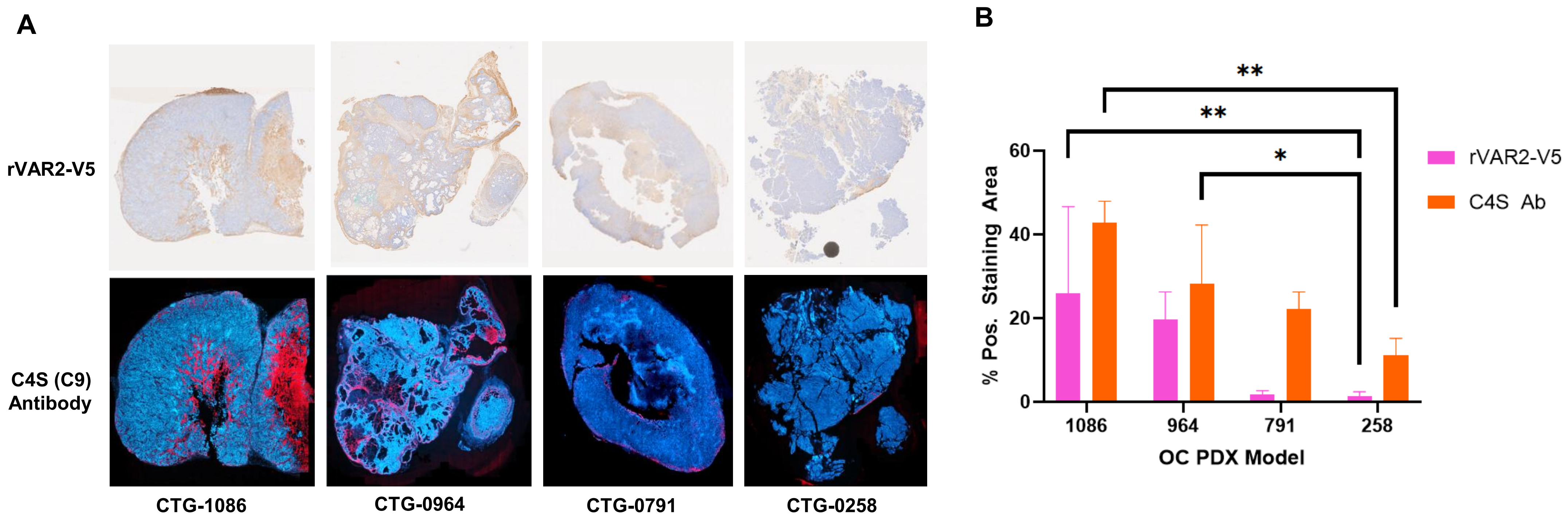


**Fig. S8 Correlation of rVAR2-V5 and C4S (C9) antibody staining. (A)** Representative ovarian cancer PDX tumors (CTG-1086, CTG-0964, CTG-0791, CTG-0258) stained with rVAR2-V5 (top row) or C4S (C9) antibody (bottom row). **(B)** Quantification of percent positive staining area calculated using QuPath. Bars represent mean ± SD of two replicate tumor sections per model. * $p < 0.05$, ** $p < 0.01$; one-way ANOVA

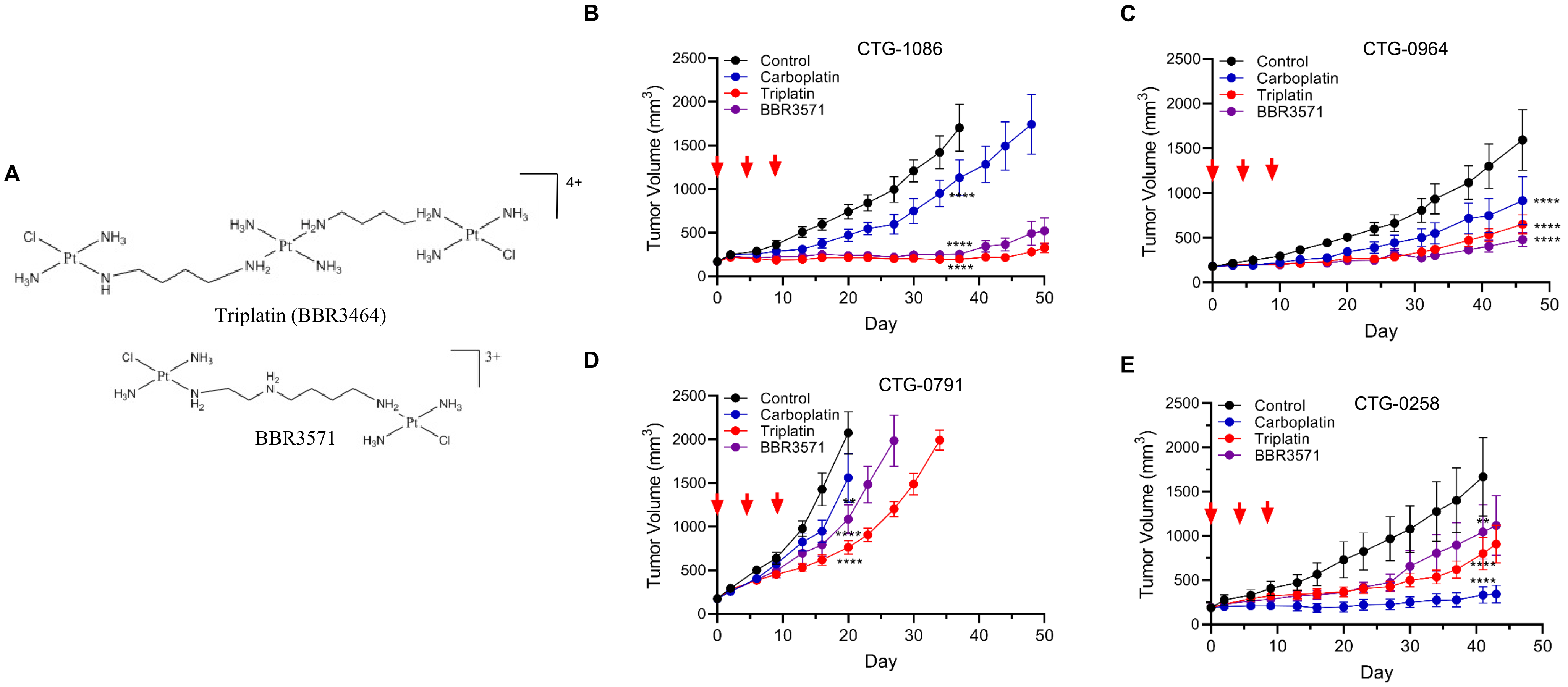


**Fig. S9. Antitumor activity BBR3571 compared to Triplatin in ovarian cancer patient-derived xenograft models. (A)** Chemical structures of Triplatin (BBR3464) and BBR3571. **(B–E)** Tumor growth in CTG-1086 (B), CTG-0964 (C), CTG-0791 (D), and CTG-0258 (E) xenografts following treatment with control (black), carboplatin (blue), Triplatin (red), or BBR3571 (purple). Red arrows indicate treatment administrations. Mice were treated i.p. with carboplatin (40 mg/kg), Triplatin (0.3 mg/kg) or BBR3571 (0.3 mg/kg) on days 0, 4 and 8. Tumor volumes are plotted against days after treatment initiation. ****$p<0.0001$, 2-way ANOVA
